# Pharmacologic Inhibition of Ferroptosis Attenuates Experimental Abdominal Aortic Aneurysm Formation

**DOI:** 10.1101/2024.06.18.599427

**Authors:** Jonathan R. Krebs, Paolo Bellotti, Jeff Arni C. Valisno, Gang Su, Shiven Sharma, Denny Joseph Manual Kollareth, Joseph B. Hartman, Aravinthan Adithan, Michael Spinosa, Manasi Kamat, Timothy Garrett, Guoshuai Cai, Ashish K. Sharma, Gilbert R. Upchurch

## Abstract

The pathogenesis of abdominal aortic aneurysm (AAA) formation involves vascular inflammation, thrombosis formation and programmed cell death leading to aortic remodeling. Recent studies have suggested that ferroptosis, an excessive iron-mediated cell death, can regulate cardiovascular diseases, including AAAs. However, the role of ferroptosis in immune cells, like macrophages, and ferroptosis-related genes in AAA formation remains to be deciphered. Single cell-RNA sequencing of human aortic tissue from AAA patients demonstrates significant differences in ferroptosis-related genes compared to control aortic tissue. Using two established murine models of AAA and aortic rupture in C57BL/6 (WT) mice, we observed that treatment with liproxstatin-1, a specific ferroptosis inhibitor, significantly attenuated aortic diameter, pro-inflammatory cytokine production, immune cell infiltration (neutrophils and macrophages), increased smooth muscle cell α-actin expression and elastic fiber disruption compared to mice treated with inactivated elastase in both pre-treatment and treatment after a small AAA had already formed. Lipidomic analysis using mass spectrometry shows a significant increase in ceramides and a decrease in intact lipid species levels in murine tissue compared to controls in the chronic AAA model on day 28. Mechanistically, *in vitro* studies demonstrate that liproxstatin-1 treatment of macrophages mitigated the crosstalk with aortic smooth muscle cells (SMCs) by downregulating MMP2 secretion. Taken together, this study demonstrates that pharmacological inhibition by liproxstatin-1 mitigates macrophage-dependent ferroptosis contributing to inhibition of aortic inflammation and remodeling during AAA formation.

## INTRODUCTION

Abdominal aortic aneurysm (AAA) is a vascular disease that causes dilation of the aorta which can lead to aortic rupture and sudden death.^1–4^ Aneurysms can occur at any location of the aorta, but the abdominal aorta is the most common site.^5^ The feared complication of AAA rupture carries a mortality rate between 65%-95% and accounts for 16,000 annual deaths.^6, 7^ One of the strongest predictors of rupture is maximum aortic diameter, with a five-year rupture rate of at least 25% for aneurysms larger than 5.0 cm.^8^ Currently, there are no directed medical therapies for AAA prevention or treatment. Therefore, there is an urgent need to develop novel targeted therapies that can attenuate aortic growth and prevent rupture.

The mechanism for AAA formation involves leukocyte recruitment and infiltration into the aortic wall media and adventitia via excessive production of pro-inflammatory cytokines and matrix degrading enzymes.^9–11^ This activates programmed cell death pathways including apoptosis and non-apoptotic pathways like neutrophil extracellular traps (NETs) and pyroptosis, that cause progressive thinning of the aortic wall, increasing the likelihood of rupture.^12–14^ The role of ferroptosis, a form of iron-dependent programmed cell death, in AAA formation has yet to be fully elucidated.^15^ Ferroptosis is triggered by oxidative damage and characterized by the accumulation of lipid peroxides in the context of increased reactive oxygen species (ROS) generation and inactivation of GSH peroxidase 4 (GPX4), a glutathione (GSH) dependent enzyme that prevents lipid peroxidation.^16–18^ It is influenced by structurally diverse small molecules (e.g. erastin, sulfasalazine, and RSL3) and also prevented by lipophilic antioxidants (liproxstatins, ferrostatins, CoQ10, vitamin E) offering a source of therapeutic potential.^19, 20^ As excessive thrombus formation and iron deposition are hallmarks of chronic AAA, an excessive iron-mediated cell death is hypothesized to immunomodulate AAA formation with a potential for therapeutic interventions using specific inhibitors of ferroptosis ^21, 22^ ^23^. We have previously shown that expression of key ferroptosis markers including increased iron content, lipid peroxidation (malondialdehyde; MDA), and depletion of GSH are present in aortic tissue of experimental murine AAAs.^24^ Furthermore, *in vitro* studies have shown that additional markers of ferroptosis including ROS production and Nrf2 nuclear translocation are significantly increased in elastase treated macrophages.^24^ Importantly, we have shown that proresolving lipid mediators such as Resolvin D1 and Maresin 1 can inhibit macrophage-dependent ferroptosis, supporting the notion that ferroptosis represents a suitable pathway for targeted pharmacologic therapy ^24^.

One of the critical hallmarks of human AAA tissue is the formation of intraluminal thrombus that is a predominant feature of proteolysis resulting in the degradation and destabilization of the aortic wall ^25^. Several components of the thrombus formation include erythrocyte trapping, hemagglutination in the thrombus, as well as ferrous iron (Fe2+), can contribute to the altered homeostasis of the vasculature. Accordingly, in this study, we evaluated the hallmarks of increased iron-mediated cell death and associated inflammatory signaling during ferroptosis, using human AAA tissue and elucidated the role of ferroptosis in murine models of chronic, thrombus forming AAA and preformed aneurysms. We hypothesize that accumulation of dead cell debris and thrombus formation leads to elevated iron and lipid peroxidation (ferroptosis), that triggers immune cell-parenchymal cell crosstalk leading to vascular remodeling, and that pharmacologic inhibition of ferroptosis attenuates ferroptosis-mediated AAA growth and prevents impending rupture.

## MATERIALS AND METHODS

### Human Single Cell RNA Sequencing

Pseudobulk analysis of all cells in a single-cell RNA sequencing dataset (GSE166676) of human AAA tissue was performed to evaluate for differentially expressed genes (DEGs) between AAA (n=4) and control (n=2).^26^ DEGs were identified via edgeR using the QL F-test and ferroptosis-related genes (FRGs) were recognized using a list of FRGs curated from Gene Cards (query=’ferroptosis’). Differential expression with absolute fold change (FC) > 2 and p-value < 0.05 was considered statistically significant.

### Animals

Adult male 8–12-week-old C57BL/6 WT mice were used in this study (Jackson Laboratory, Bar Harbor, ME). Mice were housed in a temperature-controlled room at 25°C in 12-hour light-dark cycles as per institutional animal protocols. Mice were provided drinking water and standard chow diet *ad libitum*. All Animal experiments followed protocol approved by the University of Florida’s Institutional Animal Care and Use committee (20220000546).

### Murine AAA Model Using Topical Elastase Treatment

C57BL/6 (wild-type; WT) mice were anesthetized using isoflurane and underwent exposure of the infrarenal abdominal aorta, as previously described ^24, 27^. The aorta was dissected circumferentially away from the surrounding tissues and subjected to 3 minutes of either 5 µL of peri-adventitial elastase (0.4 U/mL type 1 porcine pancreatic elastase, Sigma Aldrich, St. Louis, MO) or heat-inactivated elastase as control. Separate groups of animals were intraperitoneally (i.p.) injected with either liproxstatin-1 (10mg/kg; Cayman Chemicals), erastin (10mg/kg, 50mg/kg, or 100mg/kg; Cayman Chemicals), or vehicle control (saline) from the day of the surgery (day 0) through post-operative day 7. The selection of liproxstatin-1 dosages were based on published literature ^28^.

Aortas were harvested on day 14 after elastase application. Mice from each group were euthanized under anesthesia by overdose and exsanguination. The abdominal aorta, from below the left renal vein to the bifurcation was dissected. The external aortic adventitia diameter was measured at its maximum diameter and at the intact self-control portion just below the left renal vein using video microscopy with NIS-Elements D.5.10.01 software (Nikon SMZ-25, Melville, NY)^25^. The aortic dilation percentage was determined using (maximal AAA diameter − self-control aortic diameter)/(maximal AAA diameter) × 100%. Aortic sections were harvested and preserved in formalin for immunohistochemistry or snap-frozen in liquid nitrogen and stored at -80 °C.

### Chronic AAA and Aortic Rupture Model

In a second chronic AAA model linked to thrombus formation and aortic rupture^29^, mice were treated with 3 minutes of peri-adventitial application of either 5 µL of peri-adventitial elastase (0.4 U/mL type 1 porcine pancreatic elastase, Sigma Aldrich, St. Louis, MO) or heat-inactivated elastase as control. Mice were given drinking water containing 0.2% Beta-Aminopropionitrile (BAPN). Separate groups of mice were treated with Liproxstatin-1 (10mg/kg administrated i.p.; Cayman Chemicals) or vehicle control (saline) daily from post-operative day 14 through post-operative day 27. Animals were euthanized on day 28 and aortic tissue was harvested as described above.

### Lipidomic Analysis

Aortic samples were extracted using the Bligh-Dyer extraction method. Chromatographic separation was performed on an ultra-high-performance liquid chromatography (UHPLC) system (Dionex UHPLC; Thermo Scientific, San Jose, CA) using ACQUITY UPLC BEH C18 1.7 µm, 2.1mm x 50 mm column with ACQUITY UPLC BEH C18 VanGuard Pre-column 1.7 µm, 2.1 mm x 5 mm with a gradient program consisting of mobile phase A (60:40 acetonitrile/water) and mobile phase B (90:10 isopropanol/acetonitrile) with 10 mmol/L ammonium formation and 0.1% formic acid was used for sample analysis. The UHPLC system was coupled to a mass spectrometer (Q-Exactive Orbitrap; Thermo Scientific) to perform global lipidomic profiling in both positive and negative ionization modes through full-scan and iterative exclusion tandem mass spectrometry. For processing the data, LipidMatch Flow was utilized to manage peak identification, remove background noise, annotate, and merge results for both ion polarities. Semi-quantitative analysis was carried out using LipidMatch Normalizer.

### Cytokine assay

Using isolated protein from murine abdominal aortas, cytokine panel assay (Bio-Rad Laboratories, Hercules, CA) was performed according to manufacturer instructions. High Mobility Group Box 1 (HMGB1) was measured in cell culture supernatants using an ELISA kit, following the manufacturer’s instructions (IBL International, Hamburg, Germany).

### Histology

Aortic tissue was fixed in 4% buffered formaldehyde for 24 hours and embedded in paraffin and sectioned at 5µm. Immunostaining was performed for elastin (Verhoeff-van Gieson), neutrophils, macrophages, and smooth muscle α-actin, as previously described^29^. Antibodies for immunohistochemical staining were anti-mouse Mac2 for macrophages (1:10,000, Cedarlane Laboratories, Burlington, ON, Canada; catalog no. CL8942AP), anti-mouse neutrophils for polymorphonuclear neutrophils (PMNs) (1:10,000, AbD Serotec, Oxford, United Kingdom; catalog no. MCA771GA), anti-mouse α-smooth-muscle-actin (α-SMA, 1:1000, Sigma, St. Louis, MO; catalog no. A5691).

Histological analysis was performed on three different sections of aortic tissue from each animal and quantified using two independent observers. Representative images from different animals from each group are depicted for accuracy. Images were acquired with 20x magnification by an Olympus microscope equipped with a digital camera using NIS-Elements D.5.10.01 software (Nikon SMZ-25, Melville, NY). For grading, the positive staining area of the entire aortic tissue sample was selected and measured using integrated optical density of each section^12^.

### *In vitro* experiments

Primary F4/80+ macrophages were isolated from WT mice following the manufacturer’s instructions (Miltenyi Biotec, Germany). Primary aortic smooth muscle cells (SMCs) were purified from WT mice as previously described.^30^ Macrophages were subsequently exposed to 5 minutes of elastase treatment followed by washing the cells with PBS and replacing the media with/without Liproxstatin-1 (10nM).^24^ Lipid peroxidation (MDA; Millipore Sigma, St. Louis, MO) and glutathione (GSH; Cayman Chemicals, Ann Arbor, MI) were measured in tissue or cell culture extracts per the manufacturer’s instructions using colorimetric assay kits. Nrf2 transcription factor activation in nuclear extracts was measured using a colorimetric assay kit (Abcam, Cambridge, UK). Cultured media transfer (CMT) experiments using macrophages and SMCs were also performed as previously described.^24^ Macrophages from WT mice were grown to confluency in 6-well plates and exposed to transient elastase treatment with/without Liproxstatin-1 treatment. After 6 hours, CMT was performed to SMC cultures and matrix metalloproteinase-2 (MMP2) activity was measured at 24 hours (Luminex bead array, Millipore Sigma).

### Statistical Analysis

Statistical evaluation was performed with GraphPad Prism 10 software (La Jolla, CA, USA). A Student’s *t* test with nonparametric Mann Whitney test was used for univariate pair-wise comparisons. Among multiple comparative groups, one-way analysis of variance (ANOVA) after post hoc Tukey’s test was performed. Values are presented as mean ± standard error of mean (SEM) and a p value of <0.05 was considered significant.

## RESULTS

### Single-cell RNA-sequencing analysis reveals dysregulation of ferroptosis-related genes in AAA patients

To evaluate for evidence of ferroptosis dysregulation in human AAA, we analyzed single-cell RNA sequencing data from AAA patients and controls using a previously reported sequencing dataset (GSE1666676), and identified differences in ferroptosis-related gene (FRG) expression^31^. Analysis was performed in pseudo-bulk of all cells and the analysis identified 1,971 differentially expressed genes (DEGs) between human AAAs and controls (Figure 1A). The list of DEGs were then cross-referenced with a curated list of ferroptosis-related genes (FRGs) that detected 287 differentially expressed FRGs in AAA samples compared to controls (Supplementary Table 1). The differentially expressed FRGs includes a diverse set of genes that participate in ferroptosis-mediated oxidative stress (Figure 1B). These findings in human AAA tissue prompted the further exploration of ferroptosis and its role in preclinical models of AAA.

**Figure 1.**
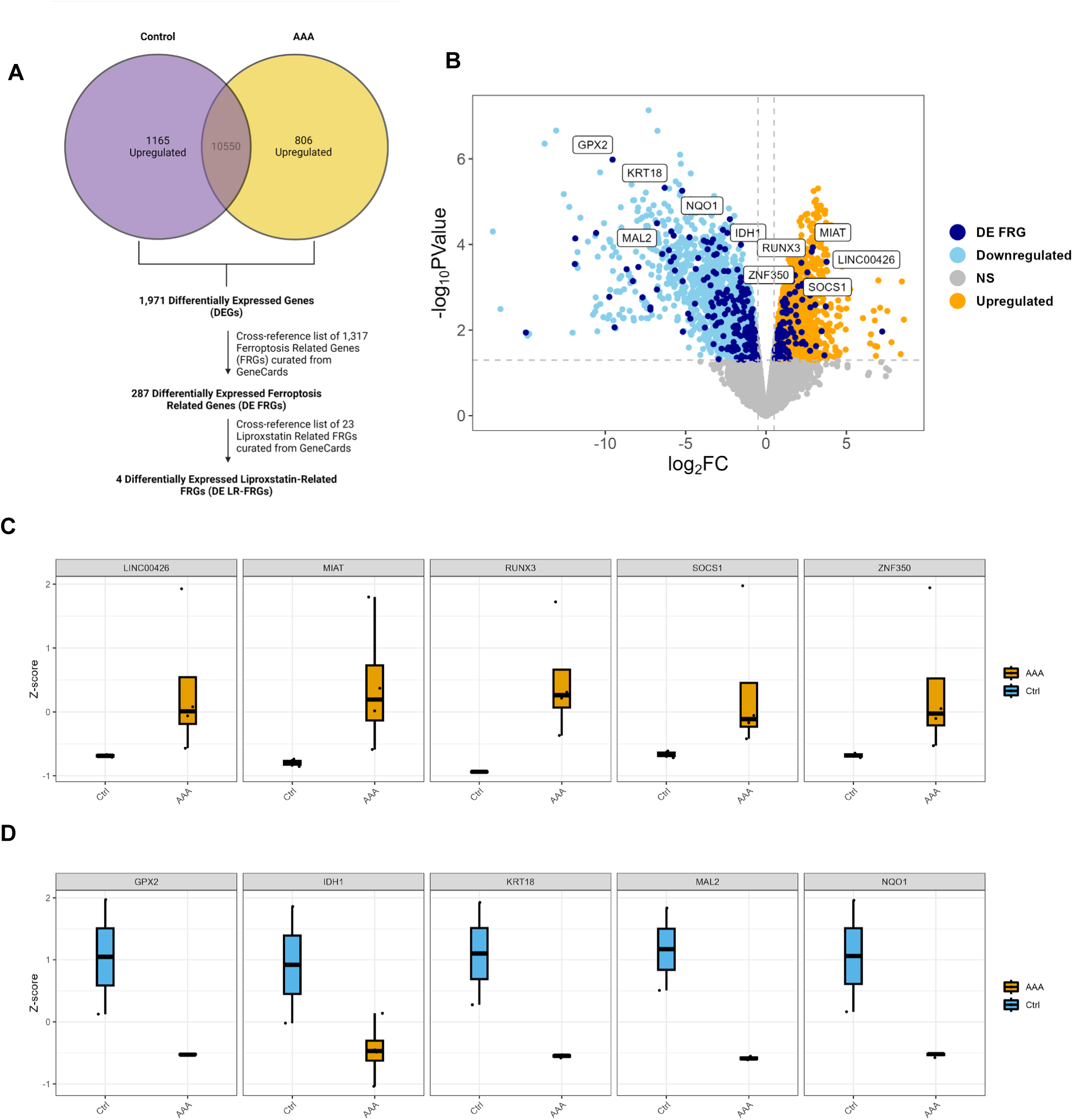
Ferroptosis-related transcripts are differentially expressed in human AAA compared to control. A, Pseudobulk analysis of publicly available single-cell RNA sequencing dataset (GSE166676). Total counts across cell populations were aggregated per sample. Differential expression analysis via edgeR revealed 1,971 differentially expressed genes (DEGs) between human AAA (n=4) and controls (n=2). A list of 1,317 ferroptosis related genes (FRGs) were curated via GeneCards search (query = ferroptosis). Cross-referencing the list with DEGs revealed 287 differentially expressed FRGs (DE FRGs). **B,** Volcano plot displaying the top 5 upregulated and downregulated genes (dark blue letters) in human AAAs compared to controls. Cut-offs for significant differential expression were set at log_2_FoldChange > 0.5 (FC > 2) and –log_10_PValue > 1.3 (p-value > 0.05). **C,** The top 5 upregulated FRGs in human AAAs: LINC00426, MIAT, RUNX3, SOCS1, ZNF350. **D,** The top 5 downregulated FRGs in human AAAs: GPX2, IDH1, KRT18, MAL2, NQO1.

### Pharmacologic inhibition of ferroptosis by Liproxstatin-1 decreases AAA formation

With this initial evidence that ferroptosis was involved in AAA, we sought to determine if pharmacologic inhibition could attenuate AAA formation in murine AAA models. First, we used the murine elastase AAA model in which aortic diameter was significantly increased in elastase-treated WT mice compared to shams (121±12.2% vs. 1.99±3.3%, p<0.0001) (Figure 2A). Administration of liproxstatin-1 significantly attenuated aortic diameter compared to untreated mice on day 14 (78.4±12.9% vs. 121±12.2%, p=0.02; Figure 2B-C). Conversely, mice were treated mice with erastin, a ferroptosis agonist, to evaluate the exacerbation of AAA formation. However, erastin treatment was not associated with increased aortic diameters on day 14 in the elastase-treated mice compared to elastase alone (Supplemental Figure S1).

**Figure 2.**
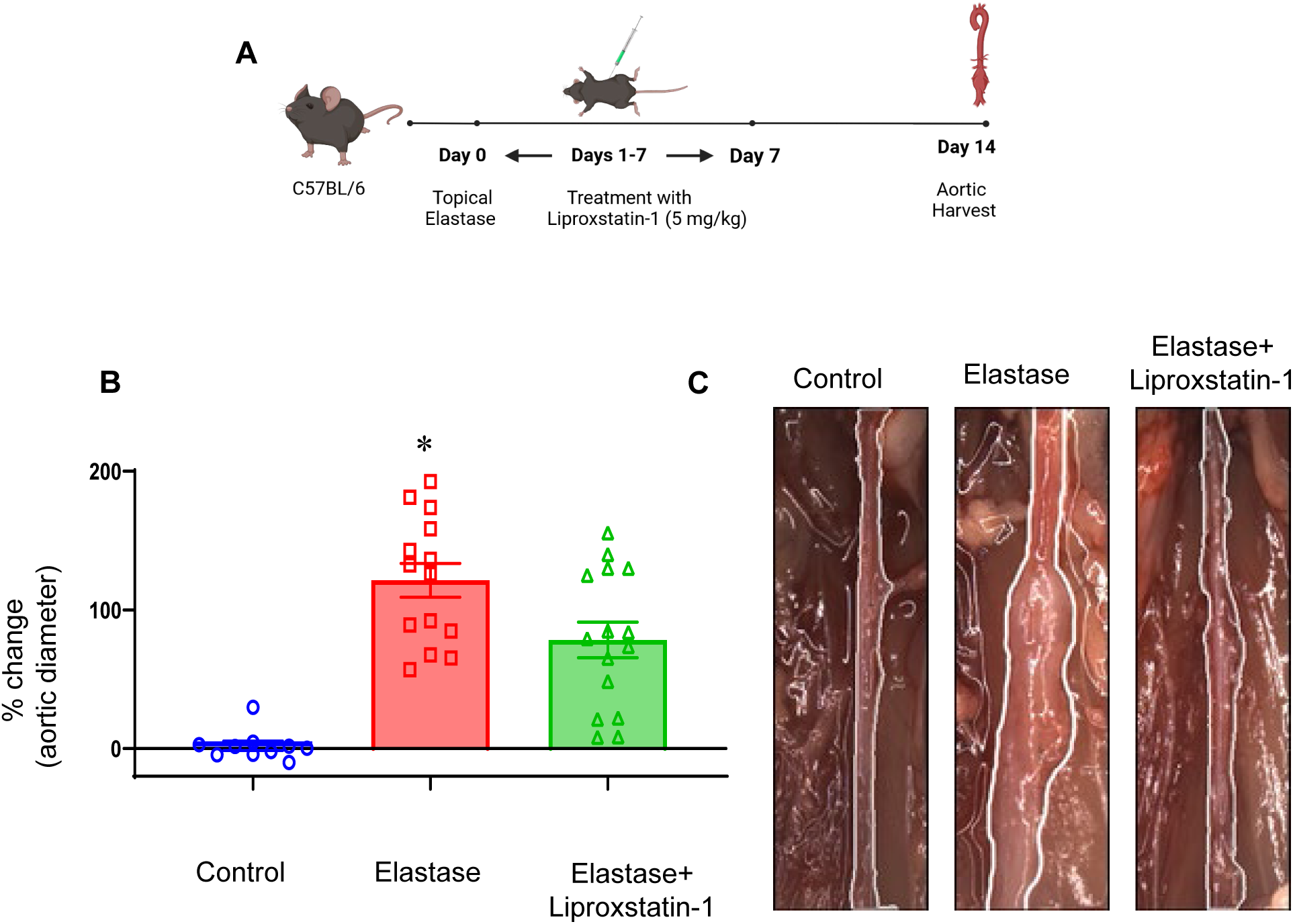
Pharmacologic inhibition using Liproxstatin-1 mitigates AAA formation in the topical elastase murine AAA model. **A,** Schematic description of the elastase treatment model of AAA. WT mice were treated with elastase or deactivated elastase (heat-inactivated; controls) with/without administration of liproxstatin-1 and aortic diameter was measured on day 14, followed by harvest of aortic tissue for further analysis. **B,** Elastase-treated mice administered with liproxstatin-1 demonstrated a significant decrease in aortic diameter compared with elastase-treated WT mice alone. *p<0.01; n=10-15/group. **C,** Representative images of aortic phenotype in all groups.

### Pharmacological inhibition of ferroptosis attenuates aortic inflammation and remodeling

Comparative histology and immunostaining of aortic tissue demonstrated distinct inflammatory cell infiltration profiles in liproxstatin-1 treated mice compared to untreated mice in the murine elastase AAA model on day 14 (Figure 3). On histological analysis elastase-treated mice demonstrated increased infiltration of neutrophils and macrophages, as well as elastin degradation, and decreased SMα-actin expression compared to sham controls. Importantly, Liproxstatin-1-treated mice demonstrated preserved aortic morphology as demonstrated by mitigation of leukocyte infiltration and elastin degradation as well as increase in smooth muscle a-actin expression compared to elastase-treated mice alone (Figure 3A-B). Additionally, a significant reduction in pro-inflammatory cytokine expression was observed in the aortic tissue of Liproxstatin-1-treated mice compared to untreated controls (Figure 4A). Moreover, aortic tissue of elastase-treated mice showed hallmarks of ferroptosis as demonstrated by a significant increase in lipid peroxidation (MDA) and decrease in GSH expression compared to sham controls, which were significantly altered in Liproxstatin-1-treated mice as observed by decrease in MDA (0.11± 0.04 vs. 0.75± 0.1 nmol/mg; p<0.01) and increase in GSH expressions (12.1±1.2 vs. 4.7±0.7 µg/mg; p=0.04) (Figure 4B-C).

**Figure 3.**
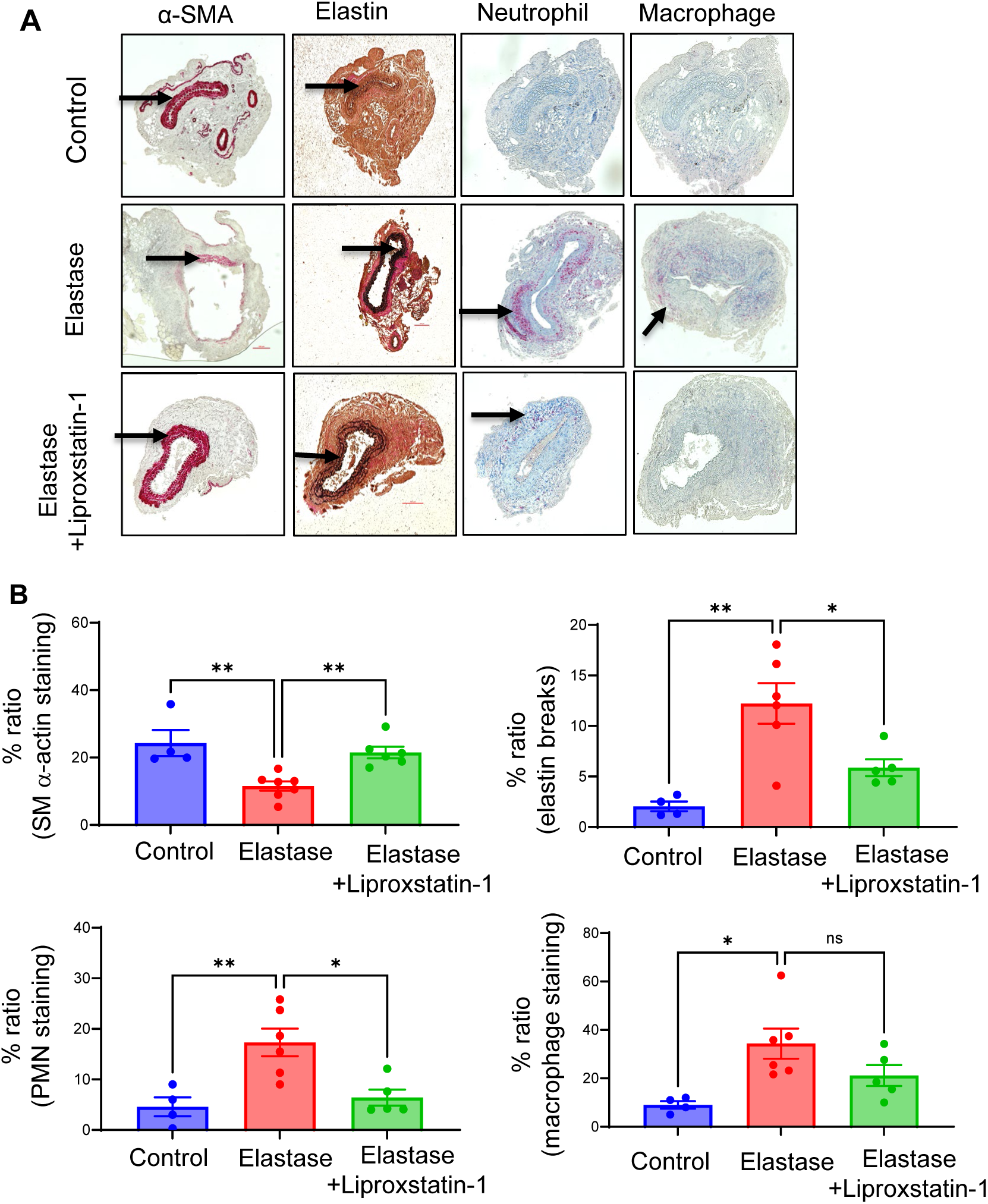
Liproxstatin-1 treatment preserves aortic morphology in the topical elastase murine AAA model. A, In liproxstatin-1 treated mice, comparative histology demonstrated increased SMα-actin expression, decreased elastin fiber disruption, and reduced immune cell infiltration (neutrophil and macrophage staining) compared to untreated mice. Arrows point to areas of immunostaining. **B**, Quantification of immunohistochemical staining demonstrates increased SMα-actin staining, and reduced elastin breaks, macrophage, and PMN staining in liproxstatin-1 treated tissue compared to mice treated with elastase alone. *p<0.05, **p<0.01; ns, not significant; n=4-7/group.

**Figure 4.**
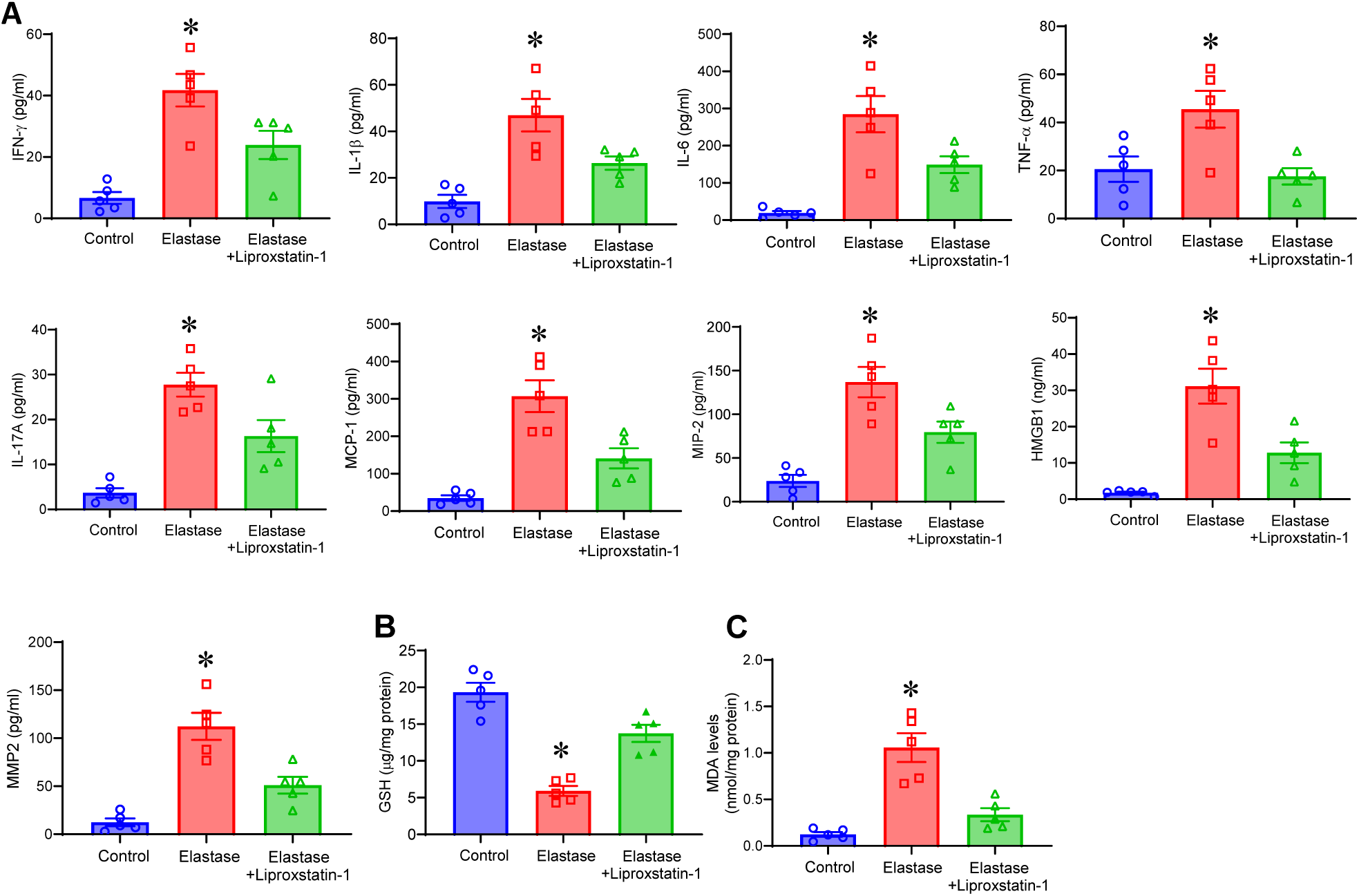
Liproxstatin-1 treatment reduces aortic inflammation and ferroptosis during AAA formation. **A**, Pro-inflammatory cytokine expression in aortic tissue of mice treated with liproxstatin-1 is significantly attenuated compared to elastase treatment alone. *p<0.01 vs. other groups; n=5/group. **B**, Glutathione depletion was significantly decreased in elastase-treated mice compared to controls. However, administration of liproxstatin-1 increased the GSH levels compared to untreated mice. *p<0.01 vs. other groups; n=5/group. **C**, Lipid peroxidation (MDA) expression in aortic tissue was significantly mitigated in liproxstatin-1 treated mice compared to elastase treatment alone. *p<0.01 vs. other groups; n=5/group.

### Inhibition of ferroptosis decreases pre-formed aneurysm growth

In order to decipher the role of ferroptosis inhibition in a pre-formed AAA, we used a second, chronic inflammatory elastase+BAPN AAA and aortic rupture model that is associated with thrombus formation.^.25^ To evaluate the protective effect of ferroptosis inhibition in a pre-formed AAA, mice were treated with Liproxstatin-1 from postoperative days 14-27, after a small aneurysm formation (Figure 5A). Mice exposed to elastase+BAPN demonstrated a significant increase in aortic diameter on day 28 compared to sham controls (507.7±69.3 vs. 3.320±0.7, p<0.0001; Figure 5B-C). More importantly, Liproxstatin-1-treated mice had significantly reduced aortic diameters compared to elastase+BAPN treated mice alone (276.4±35.9 vs. 507.7±69.; p<0.005) (Figure 5B-C).

**Figure 5.**
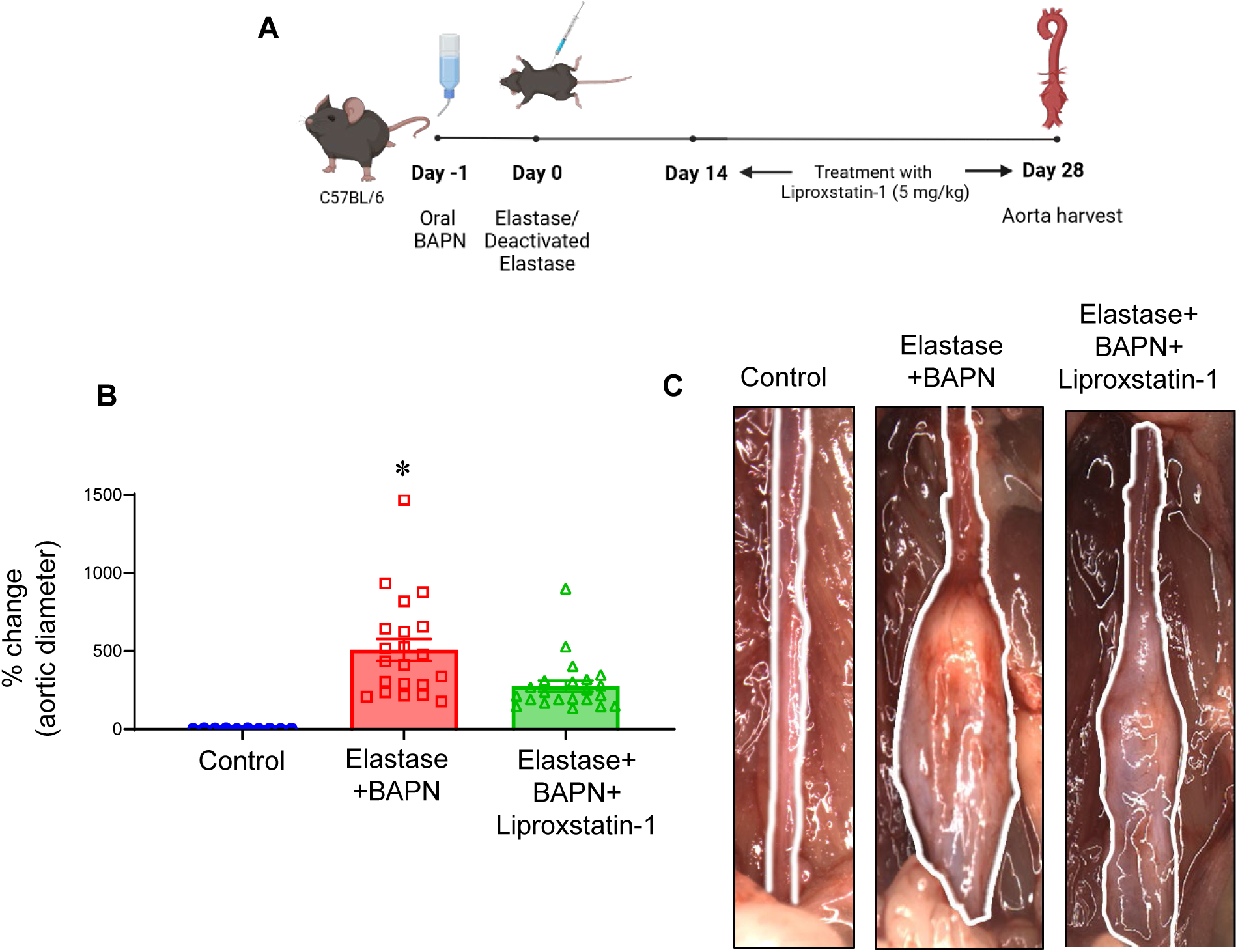
Liproxstatin-1 treatment protects against lipid oxidation during AAA formation. **A-B**, Schematic description of the elastase+BAPN treatment model of AAA with/without administration of liproxstatin-1 followed by harvest of aortic tissue for lipidomic analysis on day 28. **C-D,** Heatmap from untargeted lipidomic analysis depicting overview of lipid profiles between groups. n=6/group; Ox, oxidized; TG, triglycerides; MGDG, monogalactosyldiacylglycerol; PE, phosphatidylethanolamine; PC, phosphatidylcholine; DG, diacylglycerols; MG, monoglyceride; SM, sphingomyelin; So, sphingosine; Cer, Ceramide. **E-F,** Principal component analysis (PCA) to determine main differences in lipid profiles between experimental groups.

### Ferroptosis-specific lipids are upregulated in experimental murine AAAs

We then sought to determine if there was evidence of ferroptosis-specific lipid breakdown products in a chronic, thrombus forming murine AAA and aortic rupture model, that we have previously described (Figure 6A).^29^ In this model, there is a significant increase in AAA growth that ultimately leads to aortic rupture. Lipidomic analysis of aortic tissue on day 28 demonstrates a significant association between ferroptosis and AAA growth. We observed significant decrease in the levels of intact lipid species in AAA mice compared to sham controls (Figure 6B). Specifically, we found decreased levels of intact lipid species such as triglycerides (TG), sphingomyelin (SM) and monogalactosyldiacylglycerol (MGDG) in aortic tissue of mice with AAAs compared to controls. Furthermore, there was significant increase in the levels of sphingosine (SO), a breakdown product of SM, as well as ceramide expression, that are activated lipids known to be associated with ferroptosis^32^, in aortic tissue of AAAs. Additionally, phospholipids with higher degree of unsaturation that render the membrane more susceptible to lipid peroxidation, such as phosphatidylcholine (PC) and phosphatidylethanolamine (PE), were significantly increased in aortic tissue of mice with AAAs compared to controls (Figure 6B-C and Supplementary Table 2).

**Figure 6.**
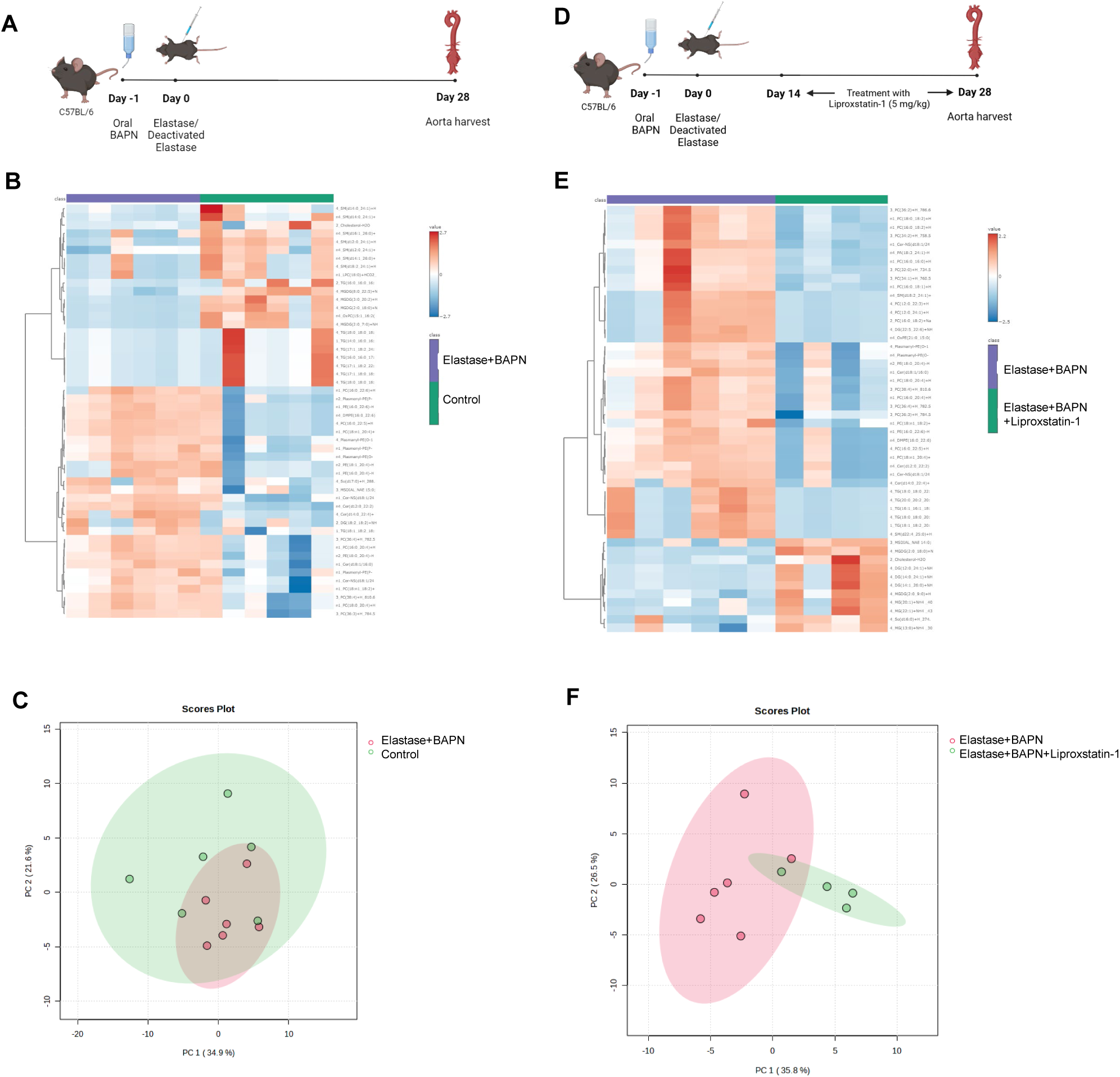
Liproxstatin-1 administration mitigates pre-formed AAAs. **A,** Schematic depicting the chronic AAA model with elastase+BAPN treatment in WT mice. Liproxstatin-1 treatment was initiated on post-operative day 14 till day 27 and analysis was performed on day 28. **B,** Aortic diameter is significantly attenuated in liproxstain-1 treated mice compared to elastase+BAPN treated mice on day 28. *p<0.002; n=10-25/group. **C,** Comparative representative of aortic phenotype in each group.

Next, we tested the effect of pharmacological inhibition of ferroptosis on the lipidomic profile in AAA mice (Figure 6D). Our results showed that Liproxstatin-1, a specific ferroptosis antagonist, offered protection against ferroptosis-related oxidized lipids in AAA. We observed a significant reduction in the levels of ceramides, PE and PC species with a higher degree of unsaturation, after Liproxstatin-1 treatment compared to untreated controls (Figure 6E-F and Supplementary Table 3). Additionally, there was a significant downregulation in the levels of oxidized PE in Liproxstatin-1 treated mice (Figure 6E). Collectively, these results indicate the involvement of ferroptosis in AAA and the efficacy of Liproxstatin-1 to attenuate ferroptosis-specific targets in the aortic tissue of mice in the chronic AAA and rupture model.

### Liproxstatin-1 treatment of pre-formed AAA mitigates aortic inflammation and aortic remodeling

Comparative histology and immunostaining demonstrated marked increase in inflammatory cell infiltration in mice treated with Liproxstatin-1 compared to untreated mice after elastase+BAPN administration. Liproxstatin-1 treated mice demonstrated significantly less neutrophil (7.12±1.2 vs. 13.2±1.9, p=0.03) and macrophage infiltrations (11.0±0.8 vs. 21.1±2.3, p=0.003), as well as increased SMα-actin expression (24.9±2.9 vs. 13.7±2.4, p=0.02) and attenuation of elastin breaks (12.8±1.6 vs. 19.0±1.8, p=0.04) compared to untreated mice (Figure 7A-B). Furthermore, aortic tissue of Liproxstatin-1 treated mice demonstrated attenuation of pro-inflammatory cytokine expression compared to untreated mice. Additionally, Liproxstatin-1 treated mice demonstrated increased GSH (1.96±0.1 vs. 0.990±0.1 nmol/mg; p=0.002), and decreased MDA (3.03±0.8 vs. 11.0±1.4; p=0.001) expressions in aortic tissue compared to elastase+BAPN-treated mice (Figure 8). These data demonstrate that Liproxstatin-1 administration can effectively inhibit ferroptosis and decrease the growth of pre-formed AAAs.

**Figure 7.**
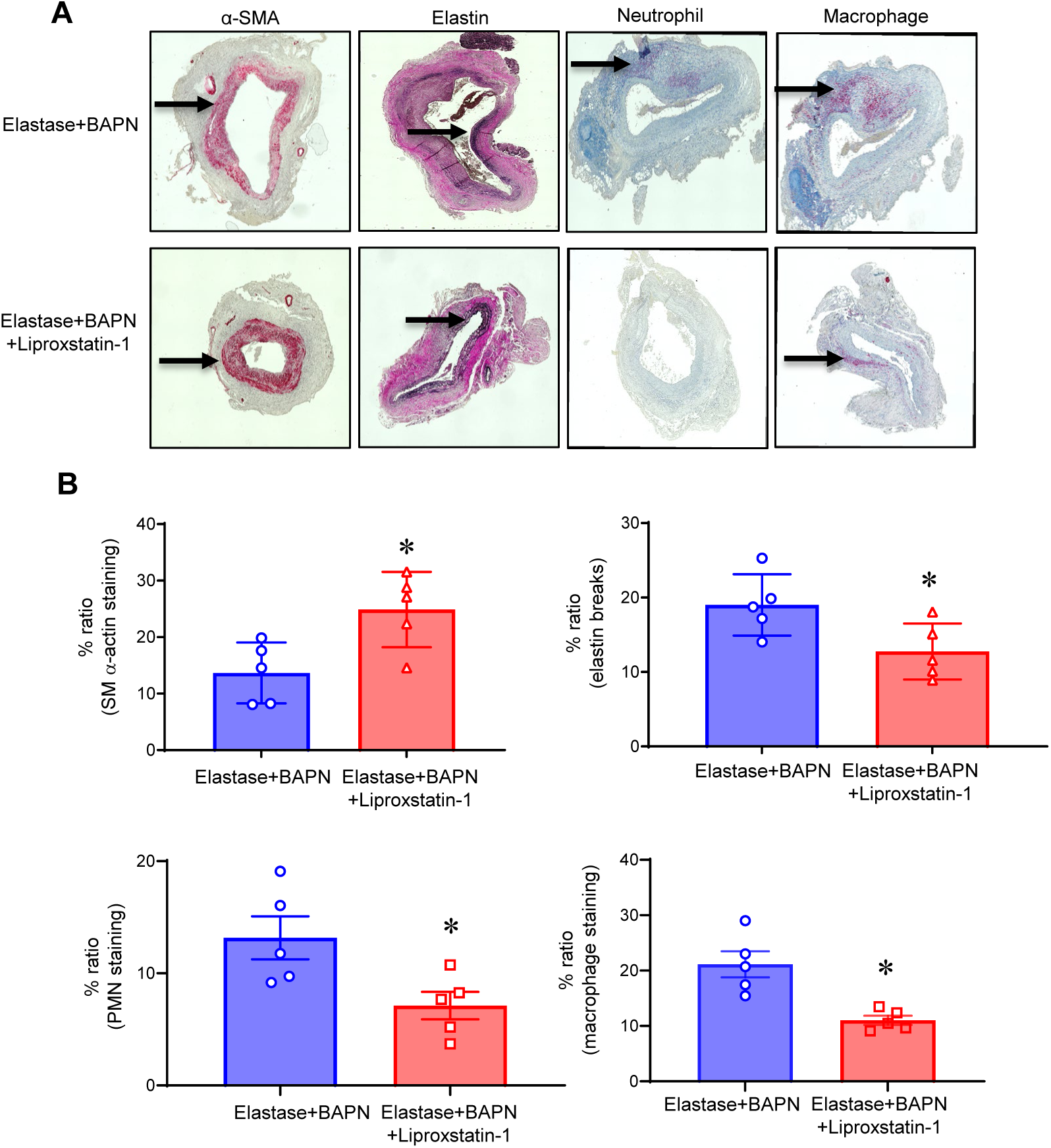
Liproxstatin-1 treatment preserves aortic morphology in the chronic murine AAA model. **A,** In liproxstatin-1 treated mice, comparative histology demonstrated increased SMα-actin expression, decreased elastin fiber disruption, and reduced immune cell infiltration (neutrophil, macrophage staining) compared to untreated mice. Arrows indicate areas of immunostaining. **B,** Quantification of immunohistochemical staining demonstrates increased SMα-actin staining, and reduced elastin breaks, macrophage, and PMN staining in liproxstatin-1 treated tissue compared to mice treated with elastase+BAPN alone. *p<0.05; n=5/group.

**Figure 8.**
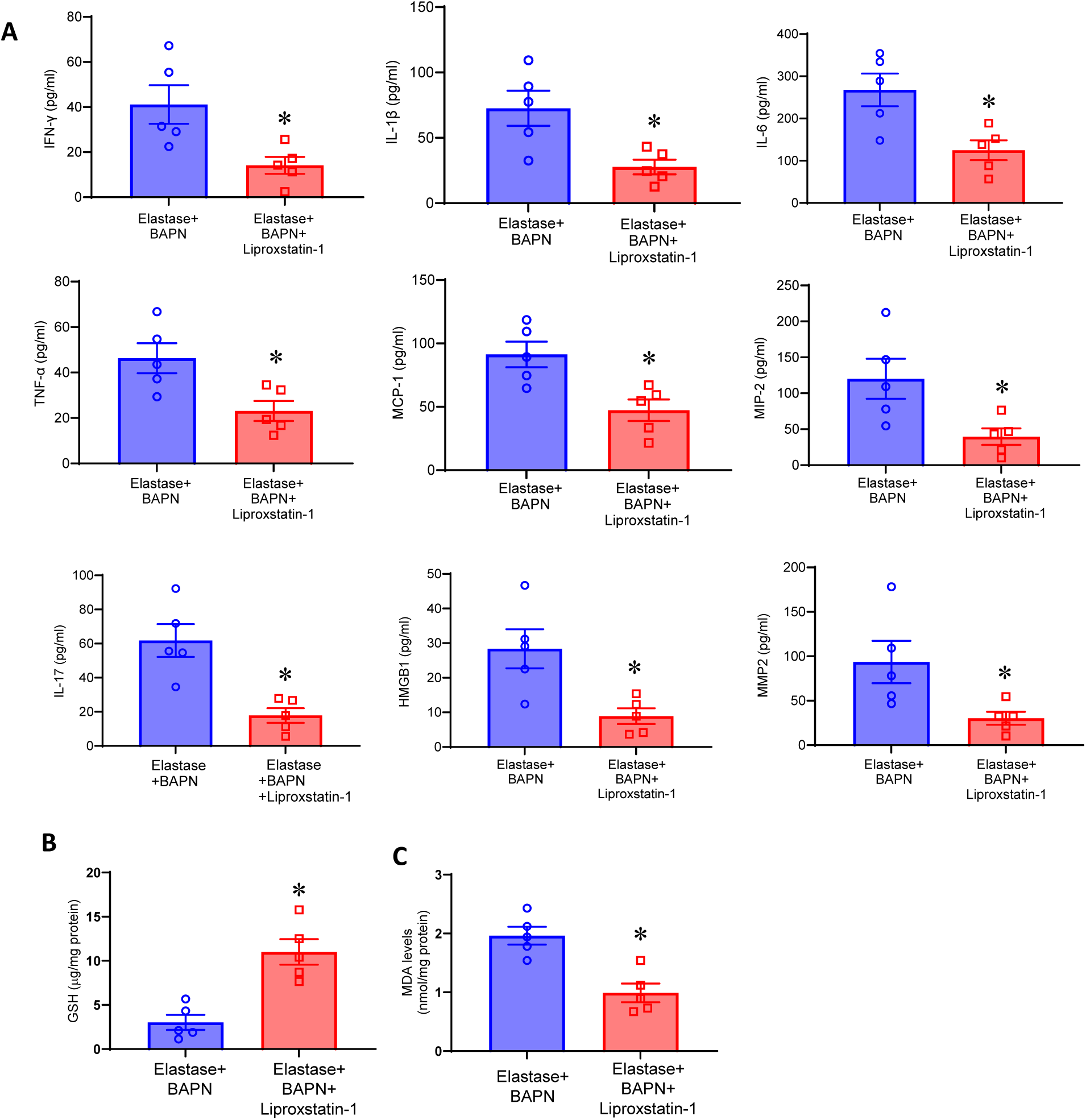
Liproxstatin-1 treatment reduces aortic inflammation and hallmarks of ferroptosis in the chronic AAA model. **A**, Aortic tissue from liproxstatin-1 administered mice in the elastase+BAPN-treated mice showed a significant decrease in pro-inflammatory cytokine/chemokine production and MMP2 expression compared to elastase+BAPN-treated WT mice. Pro-inflammatory cytokine expression in aortic tissue treated with liproxstatin-1 is significantly mitigated compared to aortic tissue treated with elastase+BAPN alone. *p<0.03, n=5/group. **B,** Elastase+BAPN treatment significantly depletes GSH levels in aortic tissue which is mitigated in mice treated with liproxstatin-1. *p<0.01; n=5/group. **C,** MDA expression in aortic tissue is significantly reduced in aortic tissue treated with liproxstatin-1 compared to untreated controls. *p<0.01; n=5/group.

### Liproxstatin-1 treatment mitigates ferroptosis in macrophages to mitigate SMC activation

To evaluate the cell-specific role of ferroptosis in AAA formation, we focused on the role of macrophages and SMCs to delineate the crosstalk between immune cell-dependent inflammation and parenchymal cell-mediated vascular remodeling in the pathogenesis s of AAA. As we had previously shown that macrophages participate as mediators of ferroptosis and histological changes were most pronounced at the smooth muscle cell predominant media layer, we hypothesized that macrophage-SMC crosstalk was important for AAA formation^24^. We first evaluated for hallmarks of ferroptosis including lipid peroxidation (MDA) and GSH depletion in macrophages exposed to transient elastase with and without Liproxstatin-1 treatment. Liproxstatin-1 treated macrophages display decreased lipid peroxidation, restoration of glutathione levels, impediment of NRF2 translocation, and reduced expression of pro-inflammatory cytokine, HMGB1 (Figure 9A-D).

**Figure 9.**
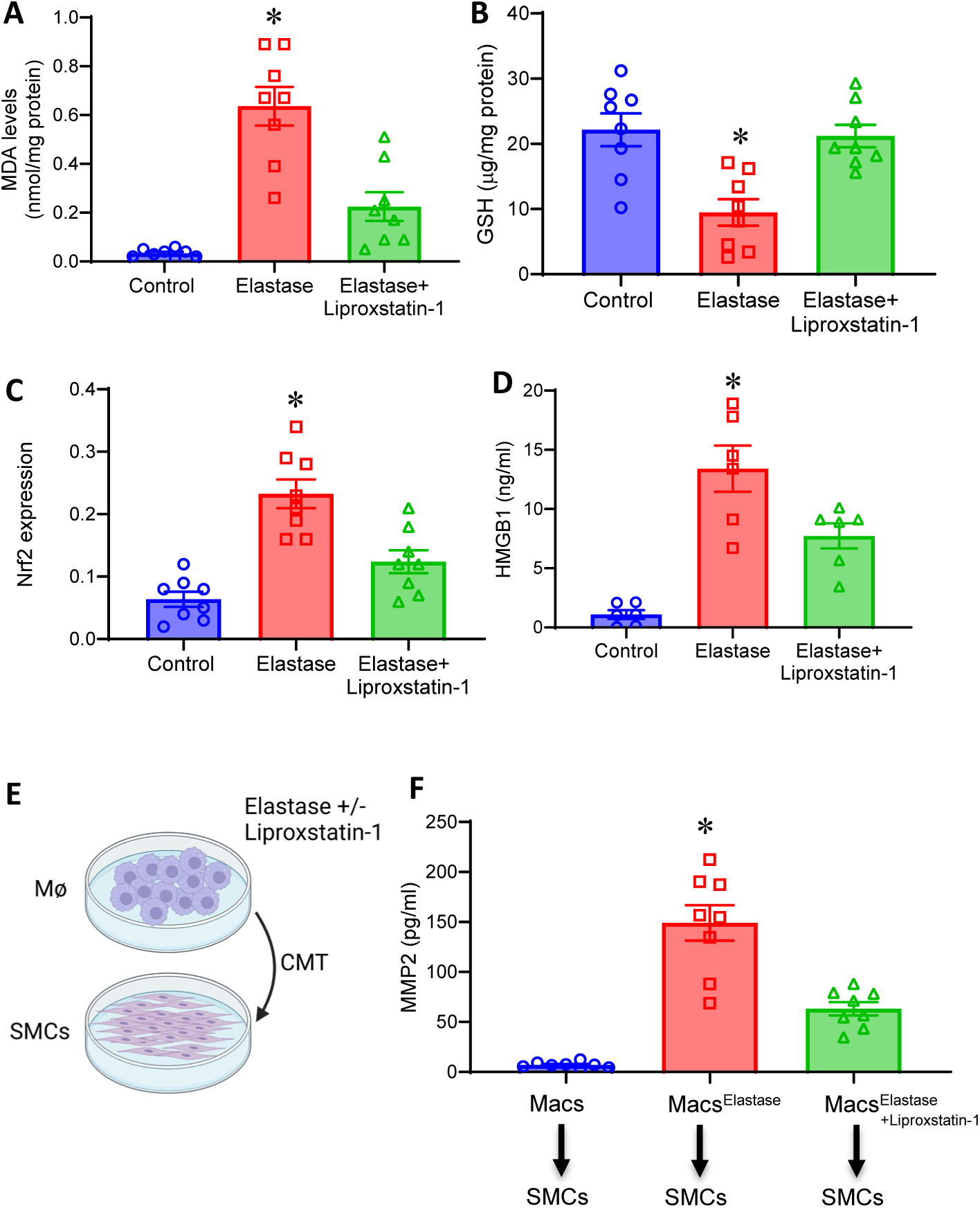
Liproxstatin-1 treatment mitigates macrophage mediated smooth muscle cell activation and MMP2 activation. **A-D,** Hallmarks of ferroptosis including increased lipid peroxidation (MDA) and glutathione depletion, as well as increased nuclear translocation of Nrf2 in cell extracts and secretion of HMGB1 in culture supernatants were observed in macrophages treated with elastase. Liproxstatin-1 treated macrophages display mitigated lipid peroxidation, restoration of glutathione levels, impediment of Nrf2 translocation, and reduced HMGB1 expression. *p<0.01 vs. other groups; n=5-10/group. **E-F,** Conditioned media transfer (CMT) from elastase-exposed macrophages induced an increase in MMP2 expression which was mitigated by liproxstatin-1 treatment. *p<0.05, **p<0.01 vs. other groups; n=5-10/group.

Next, we performed conditioned media transfer of elastase-exposed macrophages to SMCs with and without Liproxstatin-1 treatment (Figure 9E). Mitigation of MMP2 activity from SMCs was observed after CMT from Liproxstatin-1 treated macrophages to SMCs compared to untreated controls (Figure 9F). Conversely, CMT from elastase-treated SMCs to macrophages did not result in production of key inflammatory cytokines such as HMGB1 or TNF-α (data not shown). Moreover, induction of ferroptosis with erastin exacerbated elastase-exposed macrophage production of HMGB1 and TNF-α, which was significantly mitigated by liproxstatin-1 treatment (Supplementary Figure S2). These results suggest that pro-inflammatory paracrine secretions from activated macrophages secondary to ferroptosis trigger SMC remodeling. This crosstalk can be significantly mitigated by inhibition of macrophage-dependent ferroptosis by treatment with Liproxstatin-1, which then attenuates vascular remodeling and reduces the growth of AAAs.

## DISCUSSION

This study demonstrates that excess iron-mediated cell death via ferroptosis is an important signaling event that triggers crosstalk between macrophages and SMCs to alter vascular inflammation and remodeling. Single cell RNA analysis of human AAA tissue demonstrated dysregulation of several ferroptosis associated-genes providing new avenues for targeted therapy. Importantly, *in vivo* and *in vitro* studies demonstrate the ability of liproxstatin-1 to pharmacologically modulate macrophage-mediated ferroptosis and mediate the balance of oxidized and intact lipid species that are the end-products of ferroptosis resulting in decreased growth of AAAs. Collectively, these results suggest an important role of ferroptosis in AAA and the efficacy of liproxstatin-1 to mitigate AAA formation and prevent aortic rupture.

The mechanism for AAA formation involves chronic damage to the aortic wall from increased production of pro-inflammatory cytokines and matrix degrading enzymes.^9, 10^ These increased inflammatory enzymes promote cell death pathways that weaken the aortic wall and contribute to AAA rupture.^15, 24^ Ferroptosis, an iron-mediated non-apoptotic cell death pathway, has recently generated significant interest in mediating the pathogenesis of cardiovascular diseases. The mechanistic role of ferroptosis may be particularly relevant to AAA due to the association between excess iron and intraluminal thrombus that often accompanies AAA.^21, 22^ In ferroptosis, intracellular transport of iron ultimately results in the formation of a oxidized lipids that increase cellular oxidative stress and generation of reactive oxygen species.^13, 17, 18^ These lipid hydroperoxides contribute to cell death via NADPH oxidase activation, a key enzyme in redox activation, which we have previously demonstrated to be involved in macrophage-dependent activation and HMGB1 secretion, thereby providing a link to pathogenesis and growth of aneurysms.^33–35^

Recent studies have demonstrated the ability of pharmacological inhibition of ferroptosis using liproxstatin-1 to prevent BAPN-mediated aortic dissection in mice.^36^ In the context of AAA, Ferrostatin-1, another pharmacologic inhibitor of ferroptosis, has been shown to reduce AAA size through GPX4 mediated vascular smooth muscle cell (VSMC) activation in the angiotensin II murine AAA model.^37^ Liproxstatin-1 has been shown to be more effective in subverting ferroptosis due to its reaction stoichiometry to trap peroxyl radicals in lipid bilayers.^38^ Using the topical elastase model, we had previously demonstrated phenotypic changes and alterations in expression of key ferroptosis markers like MDA excess and glutathione depletion in murine aortas.^24^ Additional studies using the topical elastase model have demonstrated that PKM2-activated T lymphocyte-derived extracellular vesicles can mediate topic elastase induced AAA formation, which is then attenuated with liproxstatin-1.^39^ However, few studies have studied the role of ferroptosis inhibition in a chronic or thrombus forming AAA model.

Induction of ferroptosis can affect glutathione peroxidase causing decreased intracellular antioxidant capacity and lipid ROS accumulation, that orchestrates redox imbalance and thrombus formation. These cellular events promote degradation of the extracellular matrix as well as smooth muscle cell remodeling during the aortic growth and progression of AAAs.^40^ Glutathione metabolism can be altered by sphingolipids, a class of lipids, including ceramides, that can regulate cell proliferation, differentiation, apoptosis, and inflammation during aneurysm formation^41^. Results from our study shows altered levels of ceramides and decreased PC and PE lipid species with higher degree of unsaturation, indicating a pivotal association of ferroptosis with lipid peroxidation in the tissue during vascular aortopathies. These oxidized lipids can lead to aortic wall injury and thrombus formation, resulting in the accumulation of immune cells and promoting the release of proinflammatory cytokines, which can aggravate vascular injury forming a self-amplified loop leading to uncontrolled growth and rupture. Key immune cells that are known to be involved in AAA formation include macrophages, neutrophils, and CD4+ T cells that orchestrate an inflammatory milieu affecting the aortic wall remodeling.^42^ Since the contributory role of ferroptosis in SMCs has been previously reported, this study focused on delineating the mechanistic pathways preceding SMC activation by immune cells such as macrophages.^43^ Excess of iron-mediated sources of macrophages include phagocytosis of senescent RBCs to produce Fe^2+^, or extracellular Fe^3+^ that enters into macrophages through TFR and is reduced to Fe^2+^.^44^ This excessive iron deposition triggers ferroptosis during conditions of altered homeostasis, which then affects paracrine secretions and surrounding parenchymal cells in the aortic tissue.

Mechanistically, we provide evidence for ferroptosis-triggered macrophages to activate SMCs and treatment with liproxstatin-1 mitigating the crosstalk between macrophages and SMCs. Ferroptosis shares several commonalities between macrophages, and iron metabolism may contribute towards M1 polarization.^45, 46^ ^47^ Studies have also shown that macrophages can activate SMCs to secrete MMPs and contribute to AAA progression.^24, 48, 49^ We first demonstrated that elastase-treated macrophages exhibit increased amounts of NRF2 nuclear translocation, which is upregulated in periods of oxidative stress. The transcription factor NRF2 mediates antioxidant response element (ARE)-related genes that regulate the expression of enzymes involved in glutathione synthesis ^50^. Importantly, synthesis and function of GPx4, intracellular iron homeostasis, and lipid peroxidation can be mediated by NRF2 target genes.^51^ Modulation of NRF2 in macrophages by liproxstatin, and subsequent mitigation of MMP2 in SMCs indicates a significant correlation of these key processes during progression of AAA formation.

The clinical relevance of ferroptosis mediated pathways is highlighted by the cell-specific analysis of aortic tissue from AAA patients. A prior study using single cell RNA sequencing of 14 AAAs and 8 transplant donor control aortas evaluated differential expression of 24 ferroptosis-related genes and found dysregulation of several of these genes in the AAA neck.^52^ We instead used a dataset containing aneurysmal aortic tissue, and deciphered a larger set of ferroptosis-related genes.^31^ This single-cell RNA-sequencing analysis showed that patients with AAA disease have dysregulation of multiple ferroptosis-related genes including AIFM2, ATF3, PKM, and CASC9, which have been shown to be ferroptosis-related and associated with liproxstatin-1 mediated signaling in murine models.^36, 39, 53^ By interactions with liproxstatin-1, these genes can directly influence GPX4 activation and effect anti-oxidative activity in AAA development.

There are several limitations to consider in this study. The experimental models used to decipher the alterations in oxidized lipids were not performed in the presence of hypercholesteremia which can be an accompanying factor in clinical AAAs. Further studies deciphering the role of metabolic changes in the aortic wall relative to thrombus formation and ferroptosis in the presence of increased cholesterol using angiotensin II and ApoE^-/-^ and Ldllr^-/-^ mice should help address the immune-parenchymal cell interactions for lipidomic perturbations in vascular pathologies. Additionally, translational efforts to expand these experiments to a porcine AAA model may provide an opportunity to bridge the gap for the use of pharmacological inhibitors of ferroptosis for protection against AAA growth. Also, although erastin treatment did not exacerbate AAA growth in the murine in vivo model, the increase of erastin-induced macrophage activation denotes the ferroptosis mediated enhancement of aortic inflammation. Additional limitations include detailed evaluation of the human sequencing datasets to perform subcluster analyses comparing differential gene expression in endothelial, macrophage, and smooth muscle cells. Ongoing studies with a robust database of cell-specific analysis of ferroptosis and lipidomic related pathways in murine and human AAA samples are postulated to expand our understanding of this critical signaling pathway. Finally, in addition to liproxstatin, several compounds that mediate iron metabolism such as deferoxamine or lipid antioxidants like Vitamin E and selenium may have the potential to mediate inhibitory effects on ferroptosis and can be investigated in future studies.

In summary, we have shown that pharmacological inhibition by liproxstatin-1 mitigates macrophage-dependent ferroptosis and decreases SMC activation that contributes to inhibition of aortic inflammation and aortic remodeling during AAA formation. Our results highlight the dysregulation of several ferroptosis-related genes in human AAA, providing potential interventional molecular targets for future therapy. Further efforts should focus on elucidating the anti-ferroptotic effects of commercially available antioxidants or drugs that alter iron metabolism as targeted therapy for degenerative AAA.

## ACKNOWLEDGMENTS

This work was supported by the following National Institutes of Health grants: NIH R01 HL138931 and RO1 HL153341 (GRU and AKS), and NIGMS postgraduate training grant T32 HL160491 (JRK). We thank Michael Spinosa and Chelsea Viscardi for help with murine experiments.

## AUTHOR CONTRIBUTIONS

A.K.S. and G.R.U. designed the research; J.K., P.B., J.V., G.S., S.S., D.K., J.H., A.A., M.K. and A.K.S. performed research; J.K., M.S., M.K.,T.G., G.C., A.K.S. and G.R.U. analyzed the data; J.K. and A.K.S. wrote the manuscript with contributions from all authors; and all authors reviewed and approved the final manuscript.

## DATA AVAILABILITY STATEMENT

The data that support the findings of this study are available in the Materials and Methods section as well as in the Supplementary Material of this article.

## Abbreviations

AAA, abdominal aortic aneurysm

Nrf2, nuclear factor-erythroid factor 2-related factor 2

MDA, malondialdehyde

SMα-actin, alpha-smooth muscle cell actin

HMGB1, high mobility group box 1

## REFERENCES

1. Johnston KW, Rutherford RB, Tilson MD, Shah DM, L H and Stanley JC. Suggested standards for reporting on arterial aneurysms. Subcommittee on Reporting Standards for Arterial Aneurysms, Ad Hoc Committee on Reporting Standards. *Society for Vascular Surgery and North American Chapter*, International Society for Cardiovascular Surgery J Vasc Surg. 13:452–458.

2. Ailawadi G, Eliason JL and Upchurch GR. Current concepts in the pathogenesis of abdominal aortic aneurysm. Journal of vascular surgery. 2003;38:584–8.

3. Dimick JB, Stanley JC, Axelrod DA, Kazmers A, Henke PK, Jacobs LA, Wakefield TW, Greenfield LJ and Upchurch GR. Variation in death rate after abdominal aortic aneurysmectomy in the United States: impact of hospital volume, gender, and age. Annals of surgery. 2002;235:579–85.

4. Bengtsson H, Sonesson B and Bergqvist D. Incidence and prevalence of abdominal aortic aneurysms, estimated by necropsy studies and population screening by ultrasound. Annals of the New York Academy of Sciences. 1996;800:1–24.

5. Sakalihasan N, Limet R and Defawe OD. Abdominal aortic aneurysm. Lancet (London, England). 365:1577–89.

6. Harris LM, Faggioli GL, Fiedler R, Curl GR and Ricotta JJ. Ruptured abdominal aortic aneurysms: factors affecting mortality rates. Journal of vascular surgery. 1991;14:812–20.

7. Anjum A, von Allmen R, Greenhalgh R and Powell JT. Explaining the decrease in mortality from abdominal aortic aneurysm rupture. The British journal of surgery. 2012;99:637–45.

8. Nevitt MP, Ballard DJ and Hallett JW. Prognosis of abdominal aortic aneurysms. A population-based study. The New England journal of medicine. 1989;321:1009–14.

9. Anidjar S, Dobrin PB, Eichorst M, Graham GP and Chejfec G. Correlation of inflammatory infiltrate with the enlargement of experimental aortic aneurysms. Journal of vascular surgery. 1992;16:139–47.

10. Aziz F and Kuivaniemi H. Role of matrix metalloproteinase inhibitors in preventing abdominal aortic aneurysm. Annals of vascular surgery. 2007;21:392–401.

11. Freestone T, Turner RJ, Coady A, Higman DJ, Greenhalgh RM and Powell JT. Inflammation and matrix metalloproteinases in the enlarging abdominal aortic aneurysm. Arteriosclerosis, thrombosis, and vascular biology. 1995;15:1145–51.

12. Spinosa M, Su G, Salmon MD, Lu G, Cullen JM, Fashandi AZ, Hawkins RB, Montgomery W, Meher AK, Conte MS, Sharma AK, Ailawadi G and Upchurch GR. Resolvin D1 decreases abdominal aortic aneurysm formation by inhibiting NETosis in a mouse model. Journal of vascular surgery. 2018;68:93S–103S.

13. Li M, Wang ZW, Fang LJ, Cheng SQ, Wang X and Liu NF. Programmed cell death in atherosclerosis and vascular calcification. Cell Death and Disease. 2022;13.

14. Lu H, Sun J, Liang W, Chang Z, Rom O, Zhao Y, Zhao G, Xiong W, Wang H, Zhu T, Guo Y, Chang L, Garcia-Barrio MT, Zhang J, Chen YE and Fan Y. Cyclodextrin Prevents Abdominal Aortic Aneurysm via Activation of Vascular Smooth Muscle Cell Transcription Factor EB. Circulation. 2020;142:483–498.

15. Lee J-Y, Kim WK, Bae K-H, Lee SC and Lee E-W. Lipid Metabolism and Ferroptosis. Biology. 2021;10.

16. Liu W, Östberg N, Yalcinkaya M, Dou H, Endo-Umeda K, Tang Y, Hou X, Xiao T, Fidler TP, Abramowicz S, Yang Y-G, Soehnlein O, Tall AR and Wang N. Erythroid lineage Jak2V617F expression promotes atherosclerosis through erythrophagocytosis and macrophage ferroptosis. The Journal of clinical investigation. 2022;132.

17. Bayır H, Anthonymuthu TS, Tyurina YY, Patel SJ, Amoscato AA, Lamade AM, Yang Q, Vladimirov GK, Philpott CC and Kagan VE. Achieving Life through Death: Redox Biology of Lipid Peroxidation in Ferroptosis. Cell chemical biology. 2020;27:387–408.

18. Yu Y, Yan Y, Niu F, Wang Y, Chen X, Su G, Liu Y, Zhao X, Qian L, Liu P and Xiong Y. Ferroptosis: a cell death connecting oxidative stress, inflammation and cardiovascular diseases. Cell death discovery. 2021;7:193–193.

19. Yoshida M, Minagawa S, Araya J, Sakamoto T, Hara H, Tsubouchi K, Hosaka Y, Ichikawa A, Saito N, Kadota T, Sato N, Kurita Y, Kobayashi K, Ito S, Utsumi H, Wakui H, Numata T, Kaneko Y, Mori S, Asano H, Yamashita M, Odaka M, Morikawa T, Nakayama K, Iwamoto T, Imai H and Kuwano K. Involvement of cigarette smoke-induced epithelial cell ferroptosis in COPD pathogenesis. Nature communications. 2019;10:3145–3145.

20. Sorokin V, Vickneson K, Kofidis T, Woo CC, Lin XY, Foo R and Shanahan CM. Role of Vascular Smooth Muscle Cell Plasticity and Interactions in Vessel Wall Inflammation. Frontiers in immunology. 2020;11:599415–599415.

21. Sawada H, Hao H, Naito Y, Oboshi M, Hirotani S, Mitsuno M, Miyamoto Y, Hirota S and Masuyama T. Aortic Iron Overload with Oxidative Stress and Inflammation in Human and Murine Abdominal Aortic Aneurysm. *Arteriosclerosis*, Thrombosis, and Vascular Biology. 2015;35:1507–1514.

22. Naito Y, Tsujino T, Masuyama T and Ishihara M. Crosstalk between Iron and Arteriosclerosis. Journal of atherosclerosis and thrombosis. 2022;29:308–314.

23. Fan B-Y, Pang Y-L, Li W-X, Zhao C-X, Zhang Y, Wang X, Ning G-Z, Kong X-H, Liu C, Yao X and Feng S-Q. Liproxstatin-1 is an effective inhibitor of oligodendrocyte ferroptosis induced by inhibition of glutathione peroxidase 4. Neural regeneration research. 2021;16:561–566.

24. Filiberto AC, Ladd Z, Leroy V, Su G, Elder CT, Pruitt EY, Hensley SE, Lu G, Hartman JB, Zarrinpar A, Sharma AK and Upchurch GR. Resolution of inflammation via RvD1/FPR2 signaling mitigates Nox2 activation and ferroptosis of macrophages in experimental abdominal aortic aneurysms. FASEB journal : official publication of the Federation of American Societies for Experimental Biology. 2022;36:e22579–e22579.

25. Piechota-Polanczyk A, Jozkowicz A, Nowak W, Eilenberg W, Neumayer C, Malinski T, Huk I and Brostjan C. The Abdominal Aortic Aneurysm and Intraluminal Thrombus: Current Concepts of Development and Treatment. Front Cardiovasc Med. 2015;2:19.

26. Davis FM, Tsoi LC, Melvin WJ, denDekker A, Wasikowski R, Joshi AD, Wolf S, Obi AT, Billi AC, Xing X, Audu C, Moore BB, Kunkel SL, Daugherty A, Lu HS, Gudjonsson JE and Gallagher KA. Inhibition of macrophage histone demethylase JMJD3 protects against abdominal aortic aneurysms. The Journal of experimental medicine. 2021;218.

27. Filiberto AC, Spinosa MD, Elder CT, Su G, Leroy V, Ladd Z, Lu G, Mehaffey JH, Salmon MD, Hawkins RB, Ravichandran KS, Isakson BE, Upchurch GR, Jr. and Sharma AK. Endothelial pannexin-1 channels modulate macrophage and smooth muscle cell activation in abdominal aortic aneurysm formation. Nat Commun. 2022;13:1521.

28. Cao Y, Li Y, He C, Yan F, Li JR, Xu HZ, Zhuang JF, Zhou H, Peng YC, Fu XJ, Lu XY, Yao Y, Wei YY, Tong Y, Zhou YF and Wang L. Selective Ferroptosis Inhibitor Liproxstatin-1 Attenuates Neurological Deficits and Neuroinflammation After Subarachnoid Hemorrhage. Neurosci Bull. 2021;37:535–549.

29. Lu G, Su G, Davis JP, Schaheen B, Downs E, Roy RJ, Ailawadi G and Upchurch GR. A novel chronic advanced stage abdominal aortic aneurysm murine model. Journal of vascular surgery. 2017;66:232–242.e4.

30. Salmon M, Johnston WF, Woo A, Pope NH, Su G, Upchurch GR, Owens GK and Ailawadi G. KLF4 regulates abdominal aortic aneurysm morphology and deletion attenuates aneurysm formation. Circulation. 2013;128:S163–74.

31. Davis FM, Tsoi LC, Ma F, Wasikowski R, Moore BB, Kunkel SL, Gudjonsson JE and Gallagher KA. Single-cell Transcriptomics Reveals Dynamic Role of Smooth Muscle Cells and Enrichment of Immune Cell Subsets in Human Abdominal Aortic Aneurysms. Ann Surg. 2022;276:511–521.

32. Thayyullathil F, Cheratta AR, Alakkal A, Subburayan K, Pallichankandy S, Hannun YA and Galadari S. Acid sphingomyelinase-dependent autophagic degradation of GPX4 is critical for the execution of ferroptosis. Cell Death Dis. 2021;12:26.

33. Cameron SJ, Russell HM and Owens AP. Antithrombotic therapy in abdominal aortic aneurysm: beneficial or detrimental? Blood. 2018;132:2619–2628.

34. McCormick ML, Gavrila D and Weintraub NL. Role of oxidative stress in the pathogenesis of abdominal aortic aneurysms. Arteriosclerosis, thrombosis, and vascular biology. 2007;27:461–9.

35. Li WG, Miller FJ, Zhang HJ, Spitz DR, Oberley LW and Weintraub NL. H(2)O(2)-induced O(2) production by a non-phagocytic NAD(P)H oxidase causes oxidant injury. The Journal of biological chemistry. 2001;276:29251–6.

36. Li N, Yi X, He Y, Huo B, Chen Y, Zhang Z, Wang Q, Li Y, Zhong X, Li R, Zhu XH, Fang Z, Wei X and Jiang DS. Targeting Ferroptosis as a Novel Approach to Alleviate Aortic Dissection. Int J Biol Sci. 2022;18:4118–4134.

37. He X, Xiong Y, Liu Y, Li Y, Zhou H and Wu K. Ferrostatin-1 inhibits ferroptosis of vascular smooth muscle cells and alleviates abdominal aortic aneurysm formation through activating the SLC7A11/GPX4 axis. Faseb j. 2024;38:e23401.

38. Zilka O, Shah R, Li B, Friedmann Angeli JP, Griesser M, Conrad M and Pratt DA. On the Mechanism of Cytoprotection by Ferrostatin-1 and Liproxstatin-1 and the Role of Lipid Peroxidation in Ferroptotic Cell Death. ACS Cent Sci. 2017;3:232–243.

39. Dang G, Li T, Yang D, Yang G, Du X, Yang J, Miao Y, Han L, Ma X, Song Y, Liu B, Li X, Wang X and Feng J. T lymphocyte-derived extracellular vesicles aggravate abdominal aortic aneurysm by promoting macrophage lipid peroxidation and migration via pyruvate kinase muscle isozyme 2. Redox Biol. 2022;50:102257.

40. Ren J, Lv Y, Wu L, Chen S, Lei C, Yang D, Li F, Liu C and Zheng Y. Key ferroptosis-related genes in abdominal aortic aneurysm formation and rupture as determined by combining bioinformatics techniques. Front Cardiovasc Med. 2022;9:875434.

41. Okrzeja J, Karwowska A and Blachnio-Zabielska A. The Role of Obesity, Inflammation and Sphingolipids in the Development of an Abdominal Aortic Aneurysm. Nutrients. 2022;14.

42. Shimizu K, Mitchell RN and Libby P. Inflammation and cellular immune responses in abdominal aortic aneurysms. Arterioscler Thromb Vasc Biol. 2006;26:987–94.

43. Zhang F, Li K, Zhang W, Zhao Z, Chang F, Du J, Zhang X, Bao K, Zhang C, Shi L, Liu Z, Dai X, Chen C, Wang DW, Xian Z, Jiang H and Ai D. Ganglioside GM3 Protects Against Abdominal Aortic Aneurysm by Suppressing Ferroptosis. Circulation. 2024;149:843–859.

44. Ma J, Zhang H, Chen Y, Liu X, Tian J and Shen W. The Role of Macrophage Iron Overload and Ferroptosis in Atherosclerosis. Biomolecules. 2022;12.

45. Handa P, Thomas S, Morgan-Stevenson V, Maliken BD, Gochanour E, Boukhar S, Yeh MM and Kowdley KV. Iron alters macrophage polarization status and leads to steatohepatitis and fibrogenesis. J Leukoc Biol. 2019;105:1015–1026.

46. Yang Y, Wang Y, Guo L, Gao W, Tang TL and Yan M. Interaction between macrophages and ferroptosis. Cell Death Dis. 2022;13:355.

47. Wen Q, Liu J, Kang R, Zhou B and Tang D. The release and activity of HMGB1 in ferroptosis. Biochemical and biophysical research communications. 2019;510:278–283.

48. Longo GM, Buda SJ, Fiotta N, Xiong W, Griener T, Shapiro S and Baxter BT. MMP-12 has a role in abdominal aortic aneurysms in mice. Surgery. 2005;137:457–62.

49. Longo GM, Xiong W, Greiner TC, Zhao Y, Fiotti N and Baxter BT. Matrix metalloproteinases 2 and 9 work in concert to produce aortic aneurysms. The Journal of clinical investigation. 2002;110:625–32.

50. Wu L, Li HH, Wu Q, Miao S, Liu ZJ, Wu P and Ye DY. Lipoxin A4 Activates Nrf2 Pathway and Ameliorates Cell Damage in Cultured Cortical Astrocytes Exposed to Oxygen-Glucose Deprivation/Reperfusion Insults. J Mol Neurosci. 2015;56:848–857.

51. Song X and Long D. Nrf2 and Ferroptosis: A New Research Direction for Neurodegenerative Diseases. Front Neurosci. 2020;14:267.

52. Zhuang J, Zhu H, Cheng Z, Hu X, Yu X, Li J, Liu H, Tang P, Zhang Y, Xiong X and Deng H. PCSK9, a novel immune and ferroptosis related gene in abdominal aortic aneurysm neck. Sci Rep. 2023;13:6054.

53. Fu D, Wang C, Yu L and Yu R. Induction of ferroptosis by ATF3 elevation alleviates cisplatin resistance in gastric cancer by restraining Nrf2/Keap1/xCT signaling. Cell Mol Biol Lett. 2021;26:26.

