## Supplemental Figures for "Pharmacologic Inhibition of Ferroptosis Attenuates Experimental Abdominal Aortic Aneurysm Formation"

**Supplemental Figure S1**

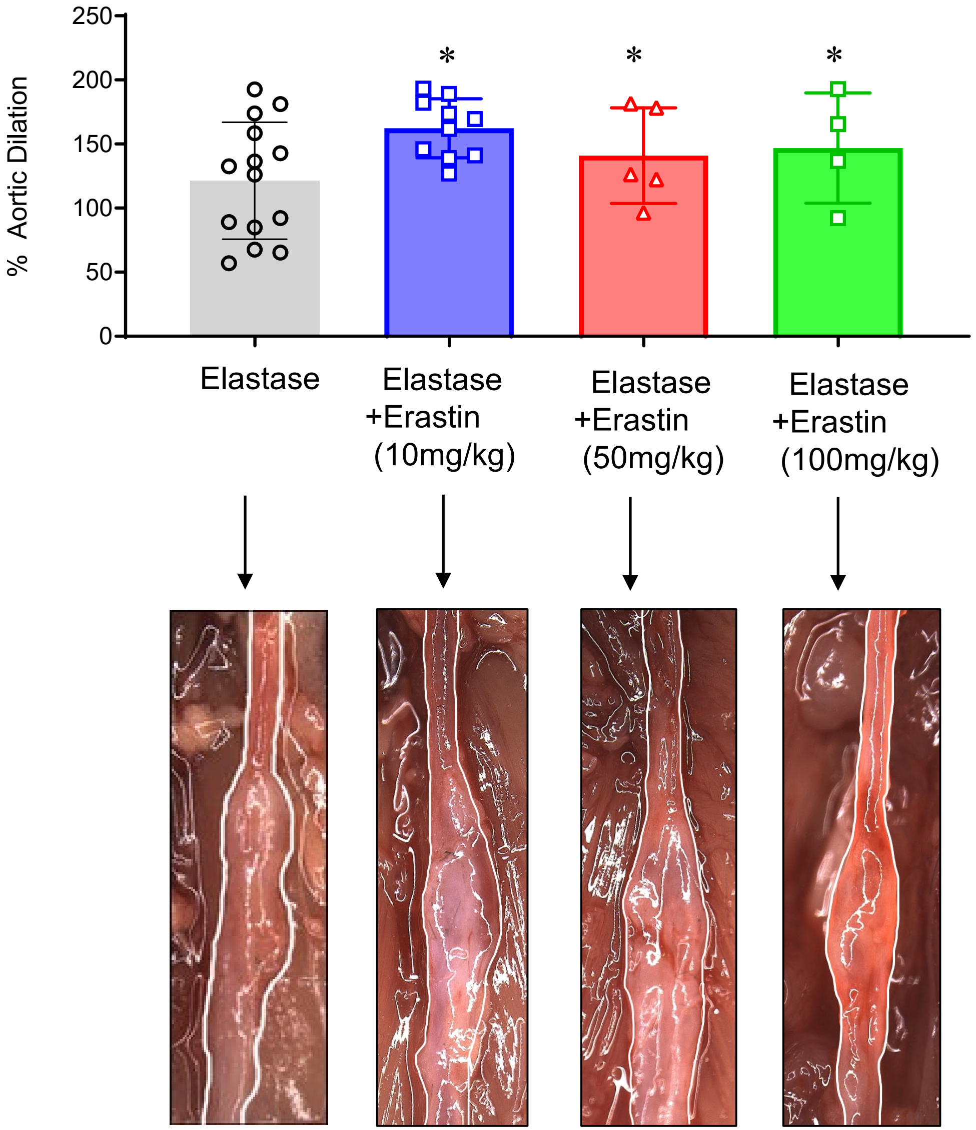


**Supplementary Figure S1.** Erastin treatment concomitantly with elastase did not exacerbate AAA growth compared to elastase alone; ns, not significant; n=5-14/group.

**Supplemental Figure S2**

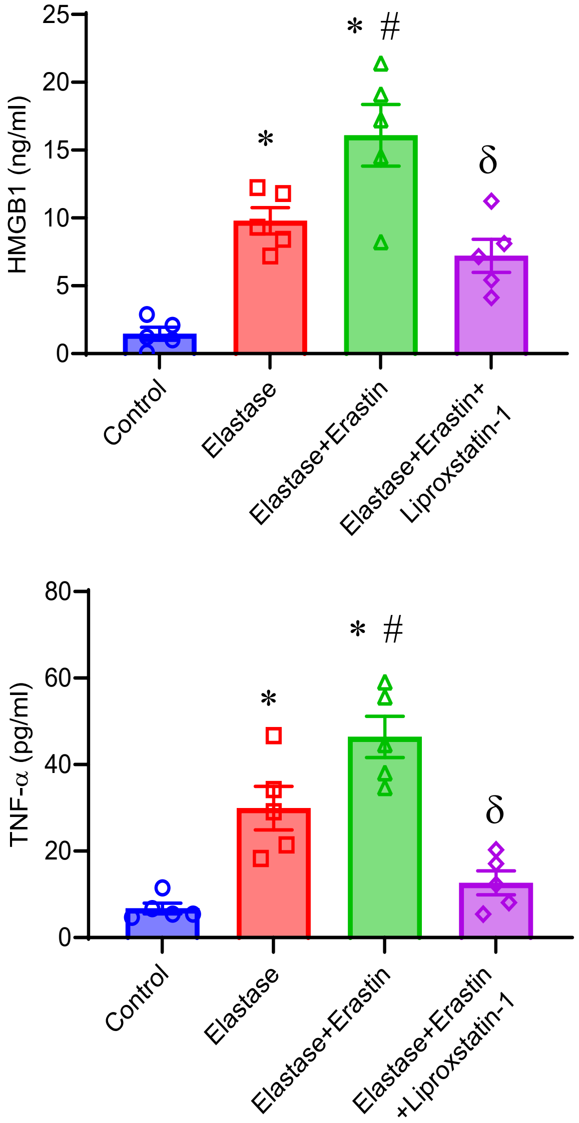


**Supplementary Figure S2.** Erastin treatment of macrophages exacerbates elastase-induced HMGB1 and TNF-α secretion that is inhibited by Liproxstatin-1. $^{*}p<0.001$ vs. control; $^{\#}p<0.02$ vs. Elastase; $^{\delta}p<0.03$ vs. Elastase+Erastin; n=5/group.
